# Implementation of a Bayesian Secondary Structure Estimation Method for the SESCA Circular Dichroism Analysis Package

**DOI:** 10.1101/2020.12.02.408302

**Authors:** Gabor Nagy, Helmut Grubmuller

## Abstract

Circular dichroism spectroscopy is a structural biology technique frequently applied to determine the secondary structure composition of soluble proteins. Our recently introduced computational analysis package SESCA aids the interpretation of protein circular dichroism spectra and enables the validation of proposed corresponding structural models. To further these aims, we present the implementation and characterization of a new Bayesian secondary structure estimation method in SESCA, termed SESCA_bayes. SESCA_bayes samples possible secondary structures using a Monte Carlo scheme, driven by the likelihood of estimated scaling errors and non-secondary-structure contributions of the measured spectrum. SESCA_bayes provides an estimated secondary structure composition and separate uncertainties on the fraction of residues in each secondary structure class. It also assists efficient model validation by providing a posterior secondary structure probability distribution based on the measured spectrum. Our presented study indicates that SESCA_bayes estimates the secondary structure composition with a significantly smaller uncertainty than its predecessor, SESCA_deconv, which is based on spectrum deconvolution. Further, the mean accuracy of the two methods in our analysis is comparable, but SESCA_bayes provides more accurate estimates for circular dichroism spectra that contain considerable non-SS contributions.

**PROGRAM SUMMARY:** Program Title: SESCA_bayes

CPC Library link to program files: (to be added by Technical Editor)

Developer’s repository link: https://www.mpibpc.mpg.de/sesca

Code Ocean capsule: (to be added by Technical Editor)

Licensing provisions: GPLv3

Programming language: Python

Supplementary material:

Nature of problem:

The circular dichroism spectrum of a protein is strongly correlated with its secondary structure composition. However, determining the secondary structure from a spectrum is hindered by non-secondary structure contributions and by scaling errors due the uncertainty of the protein concentration. If not taken properly into account, these experimental factors can cause considerable errors when conventional secondary-structure estimation methods are used. Because these errors combine with errors of the proposed structural model in a non-additive fashion, it is difficult to assess how much uncertainty the experimental factors introduce to model validation approaches based on circular dichroism spectra.

**Solution method:** For a given measured circular dichroism spectrum, the SESCA_bayes algorithm applies Bayesian statistics to account for scaling errors and non-secondary structure contributions and to determine the conditional secondary structure probability distribution. This approach relies on fast spectrum predictions based on empirical basis spectrum sets and joint probability distribution maps for scaling factors and non-secondary structure distributions. Because SESCA_bayes estimates the most probable secondary structure composition based on a probability-weighted sample distribution, it avoids the typical fitting errors that occur during conventional spectrum deconvolution methods. It also estimates the uncertainty of circular dichroism based model validation more accurately than previous methods of the SESCA analysis package.

## 1. Introduction

Circular dichroism (CD) spectroscopy in the far ultraviolet (UV) range (175-260 nm) is an established method to study the structure of proteins in solution [1, 2], because of the conformation-dependent characteristic CD signal of peptide bonds that comprise the backbone of all proteins and oligopeptides. In particular, the CD spectrum is known to change with the secondary structure (SS) of proteins, and markedly different spectra are observed for proteins rich in *α*-helices, *β*-sheets, and disordered regions [3, 4]. Because of these characteristic signals, it is common to interpret CD spectra by decomposing them into a set of basis spectra that each represent the average CD signal of pure (secondary) structure elements.

The CD analysis package SESCA (Structure-based Empirical Spectrum Calculation Approach) [5] allows for using several empirical basis spectrum sets in two methods. The first method predicts a theoretical CD spectrum from a proposed SS composition, which is typically obtained from a model structure or structural ensemble. The second method fits a measured CD spectrum to estimate the protein SS composition. Both methods can be used to validate protein structural models. The accuracy and precision of validation methods is mainly limited by scaling errors due to the uncertainty of the measured protein concentration and non-SS contributions that are not represented in the basis spectra. We have quantified the uncertainty caused by these deviations between measured CD spectra and their predicted SS signals previously [6].

The same study also revealed a potential caveat in the current SS estimation method used in SESCA. In this deconvolution method, a linear combination of selected basis spectra is used to approximate a measured CD spectrum of the protein of interest. The coefficients of the approximation with the smallest deviation are used to estimate the fraction of SS elements in the protein under the measurement conditions. Unfortunately, the interference caused by non-SS contributions may increase the deviation from the measured spectrum for some SS compositions and decrease it for others, which may lead to significant errors in deconvolution-based SS estimates.

To alleviate this problem, we developed and implemented a new SS estimation method for SESCA. The Python module, SESCA_bayes determines the likelihood of putative SS compositions using a Bayesian inference framework for a given measured CD spectrum and a basis spectrum set. This method uses the expected joint probability distribution of deviations caused by scaling errors and non-SS contributions, and thus fully accounts for the uncertainty caused by these two experimental factors. Here, we describe the theoretical background, general workflow, as well as input and output parameters of this implementation. Further, we will assess the accuracy and precision of this method through a series of sample applications.

## 2. Theory: Bayesian SS probabilities

Our goal using this method is to determine the conditional probability *P* (*SS|CD*) of SS compositions given a previously measured CD spectrum. According to Bayes’ rule [7], this probability can be inferred according to

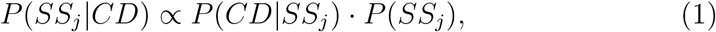

where *P* (*CD|SS_j_*) is the probability of observing the measured spectrum for a protein with a given SS composition *j* (*i.e*, the likelihood function) and *P* (*SS_j_*) is the prior probability of the given SS composition of the protein.

As shown in Fig. 1 (top), the likelihood *P* (*CD|SS_j_*) is determined in five steps. First, the SS signal is predicted from the SS composition of interest (*C_ji_*) using an appropriate basis spectrum set (*B_il_*), as discussed in our previous study [5]. Second, if the basis set provides side chain corrections based on the protein sequence, they are added to the predicted spectrum. Third, the measured CD spectrum is rescaled to minimize the root-mean-square deviation (RMSD) from the predicted spectrum using

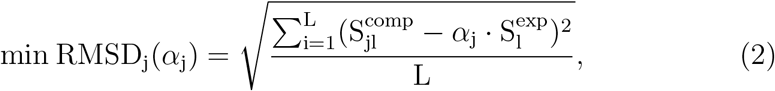

where 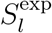 and 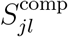 are CD intensities of the measured CD spectrum and the spectrum computed for *SS_j_* at wavelength *l*, respectively. The obtained scaling factor *α_j_* quantifies and eliminates deviations from scaling errors of the measured spectrum, whereas the RMSD from the rescaled spectrum (*RMSD_j_*) quantifies the average deviation due to unaccounted non-SS contributions. Once *RMSD_j_* and *α_j_* (collectively CD deviations) are computed, the likelihood of such deviations is determined from the joint probability distribution (*P_RMSD,α_*, see below) to estimate the likelihood of observing the measured CD spectrum for the given SS composition *P* (*CD|SS_j_*). Finally, to compute the posterior probability *P* (*SS_j_|CD*) of SS composition *j*, the CD spectrum likelihood is multiplied by the prior SS probability.

**Figure 1:**
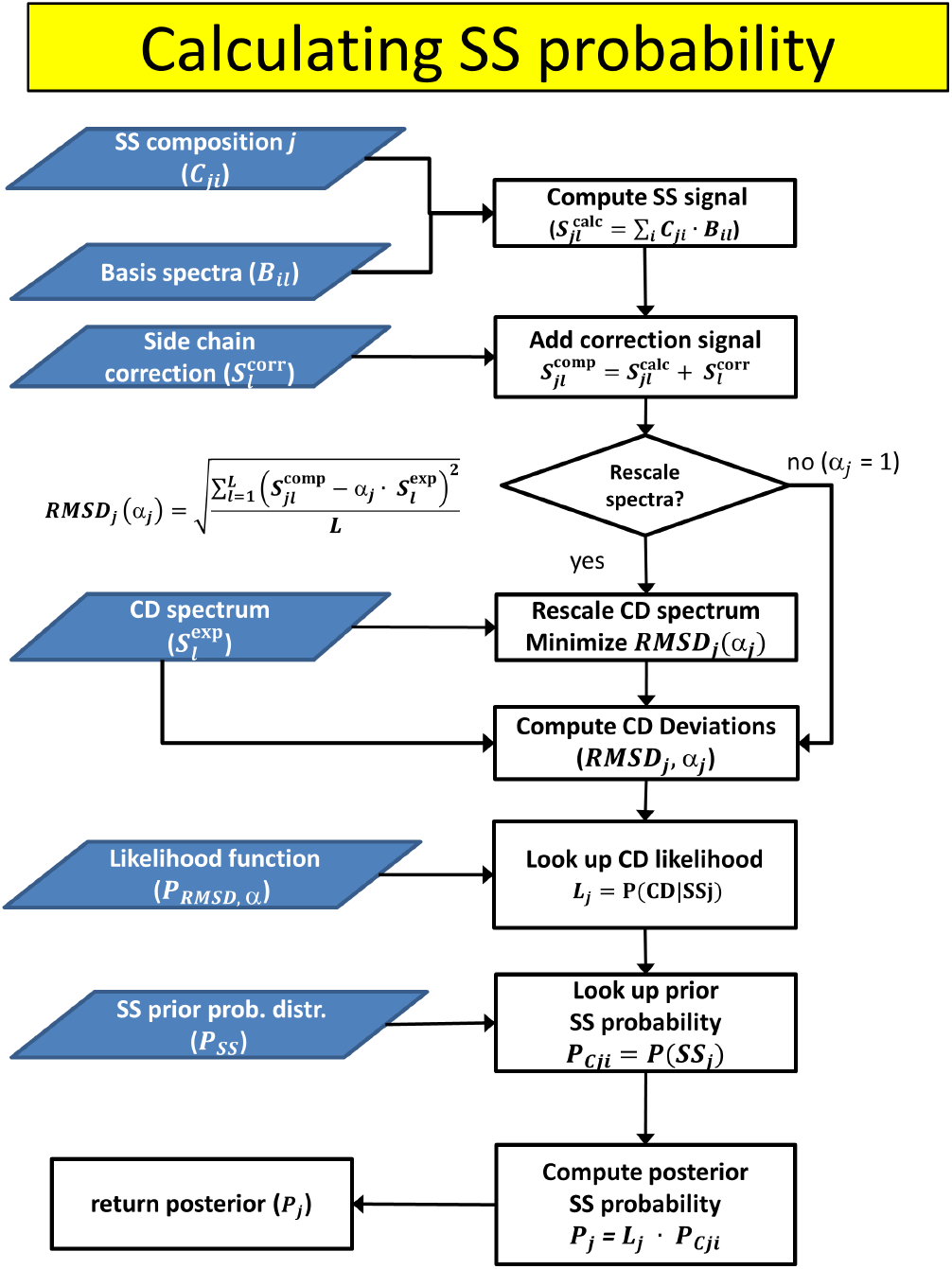
Secondary structure probability calculation scheme. The figure depicts the algorithm to compute the posterior probability of a given secondary structure *j*, based on its prior probability, and the deviations between its predicted CD signal and a given measured CD spectrum. Input data are depicted as blue parallelograms, operations as white rectangles, and decisions as white diamonds.

## 3. Methods

### 3.1. Joint probability distributions

We computed discrete joint-probability distribution functions for SESCA_bayes that can be used to determine CD spectrum likelihoods. These probability distributions were computed from CD deviations extracted from SS estimations of previously measured CD spectra. Reference CD spectra were taken from the SP175 reference set [8], which contains 71 synchrotron radiation CD (SR-CD) spectra of globular proteins with varying SS compositions. The CD spectrum of Jacalin (SP175/41) was discarded from the data set due to issues reported during the measurement and its unusually large estimated CD deviations.

The joint probability distribution functions of CD deviations were constructed as the sum of 70 two-dimensional Gaussian functions, each representing the estimated scaling factors and non-SS contributions of a reference spectrum from the SP175 set. The mean and the variance of these Gaussian functions was determined by averaging over multiple *RMSD_j_* and *α_j_* values obtained for each CD spectrum from SS estimations using four different basis spectrum sets. This approach yielded likelihood functions that were defined for a wide range of possible CD deviations, and took the uncertainty due to discretization errors of the basis spectrum determination into account.

In SESCA there are two types of basis sets, those that are solely based on SS compositions, and those that also include side chain corrections. Because the average size of CD deviations differs for these two basis set types, we determined two probability distributions shown in Fig. 2. The joint probability distribution function for basis set without side-chain corrections (left) was calculated from CD deviations estimated using the basis sets DS-dT, DSSP-1, HBSS-3, and DS5-4. For basis sets including side-chain corrections, the joint probability of CD deviations (right) were computed using the basis sets DS-dTSC3, DSSP-1SC3, HBSS-3SC1, and DS5-4SC1. For clarity, the Figure shows both a linear (top row) as well as logarithmic (bottom row) representation of the CD deviation likelihood. For both likelihoods, the onedimensional probability distribution of *RMSD_j_* was also calculated, which can be used to estimate the secondary structure from CD spectra without regards to the applied scaling factors, albeit these estimates naturally have a lower precision.

**Figure 2:**
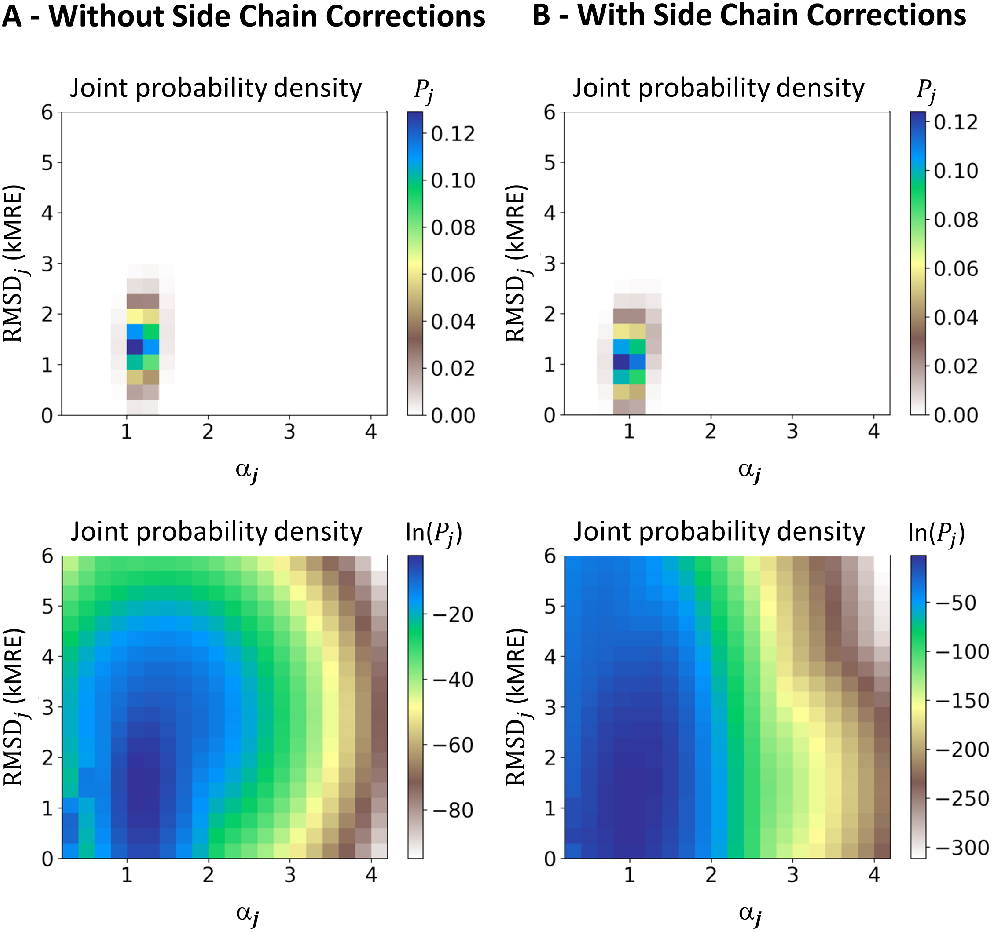
The panels depict the heat map representation of two likelihood functions provided for Bayesian SS estimation with SESCA. The estimated joint-probability distributions are shown for basis spectra that A) predict CD signals solely from SS information (left) and B) also include CD corrections from sequence-based side-chain information (right). Panels on the top and bottom show the same probability distributions using a linear and logarithmic color scale, respectively.

### 3.2. Synthetic spectra

To test the accuracy of the Bayesian SS estimation method, eight synthetic CD spectra were created using a linear combination of the three basis spectra from the DS-dT basis set (as discussed in our previous study [5]). To this aim, the coefficients shown in Table 1 for the basis spectra *α*-helix, *β*-strand, and Other for each spectrum were used. For five of eight synthetic spectra (k= 1 to 5), random coefficients were generated from uniformly distributed random numbers between zero and one, subsequently normalized to sum up to one. For the sixth synthetic spectrum (k= 6), the coefficients 0.3, 0.4, and 0.3 as well as the non-SS contributions (see below) were adopted from our previous study [6] for comparison. Synthetic spectra seven and eight were generated using low *α*-helix and *β*-strand contents respectively to represent intrinsically disordered proteins (IDP).

**Table 1:** SS compositions and CD deviations of model proteins. Columns show the index and name of the respective model protein, the fraction of its amino acids assigned to the SS classes *α*-helix, *β*-strand, and Other, as well as scaling factors *α_k_* and root-mean squared intensities *RMSI_k_* of non-SS signals to quantify scaling errors and non-SS contributions in the protein CD spectrum, respectively. Synth denotes synthetic spectrum in the proteins name, whereas Lysm, Dqd-1, Sub-C, and MeV-Nt abbreviate Lysozyme, Dehydroquinate dehydratase I, Subtilisin Carlsberg, Measles Virus Nucleoprotein C-terminal domain respectively. Note that SS fractions, scaling factors, and non-SS contributions for all synthetic proteins (k= 1-8) were parameters used to generate their CD spectrum, whereas for reference proteins (k= 9-12), all values were computed based on their measured spectra and protein data bank structures (193L, 2DHQ, 1KU8, and 1SCD, respectively).For the disordered Mev-Nt, the reference parameters were determined from the CD spectrum by Longhi *et al.* and a molecular dynamics ensemble by Robustelli *et al.* [9]

| k | protein | α-helix | β-strand | Other | $\alpha_k$ | $RMSI_k$ |
| --- | --- | --- | --- | --- | --- | --- |
| 1 | Synth-1 | 0.11 | 0.40 | 0.49 | 1.0 | 0.0 |
| 2 | Synth-2 | 0.41 | 0.20 | 0.39 | 1.0 | 0.4 |
| 3 | Synth-3 | 0.43 | 0.10 | 0.47 | 1.2 | 2.0 |
| 4 | Synth-4 | 0.27 | 0.26 | 0.47 | 1.6 | 2.7 |
| 5 | Synth-5 | 0.00 | 0.33 | 0.67 | 1.4 | 0.7 |
| 6 | Synth-6 | 0.30 | 0.40 | 0.30 | 1.0 | 3.6 |
| 7 | Synth-7 | 0.11 | 0.00 | 0.89 | 1.1 | 2.3 |
| 8 | Synth-8 | 0.00 | 0.08 | 0.92 | 1.3 | 1.3 |
| 9 | Lysm | 0.35 | 0.03 | 0.62 | 1.1 | 1.0 |
| 10 | Dqd-1 | 0.43 | 0.18 | 0.39 | 1.1 | 2.9 |
| 11 | Jacalin | 0.01 | 0.28 | 0.71 | 0.3 | 3.2 |
| 12 | Sub-C | 0.30 | 0.12 | 0.58 | 0.4 | 1.2 |
| 13 | MeV-Nt | 0.08 | 0.03 | 0.89 | 1.7 | 1.8 |

To model the effects of experimental deviations from the ideal SS signal, the CD spectra were modified by adding non-SS signals and scaling errors. The size of these CD deviations for each synthetic spectrum was quantified by the scaling factors *α_k_* and the root-mean-squared intensities of non-SS signals *RMSI_k_* listed in Table 1. Synthetic spectrum 1 (k= 1) was a positive control without any CD deviations (*α_k_*= 1.0, *RMSDI_k_*= 0.0 kMRE), spectra 2 and 6 included small (0.4 kMRE) and large (3.5 kMRE) non-SS deviations, respectively, but no scaling errors. CD deviations for spectra 3, 4, 5, 7 and 8 were drawn from the marginal distributions of experimentally observed scaling factors and non-SS contributions using rejection sampling.

Non-SS signals were generated as sums of Gaussian functions using

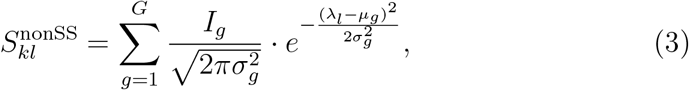

where the non-SS signal 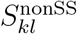 of spectrum *k* at wavelengths *λ_l_* from 178 to 269 nm was computed from the following randomly chosen parameters. The number of Gaussians *G* was chosen from the range 1 to 5, the relative peak intensity for Gaussian *g I_g_* was chosen between −20.0 and 20.0, with a peak position *μ_g_* chosen from 178 to 241 nm, and peak half-widths *σ_g_* chosen between 2 and 37 nm. Once the parameters were determined, the non-SS signal at every wavelength (using 1 nm spacing) was calculated, and the non-SS signal intensity was rescaled to match the previously defined RMSI values in Table 1.

The final synthetic spectra were computed by determining the SS signals first, by adding the appropriately scaled non-SS signal contributions in a second set, and finally by rescaling the resulting CD spectrum according to the indicated scaling factor.

## 4. Algorithm overview

Our newly implemented Python module SESCA_bayes.py performs a Monte-Carlo (MC) sampling in SS space to determine the most probable SS composition of a protein based on its measured CD spectrum. Figure 3 shows the flowchart of the algorithm that is divided into three phases: preparation, sampling, and evaluation.

**Figure 3:**
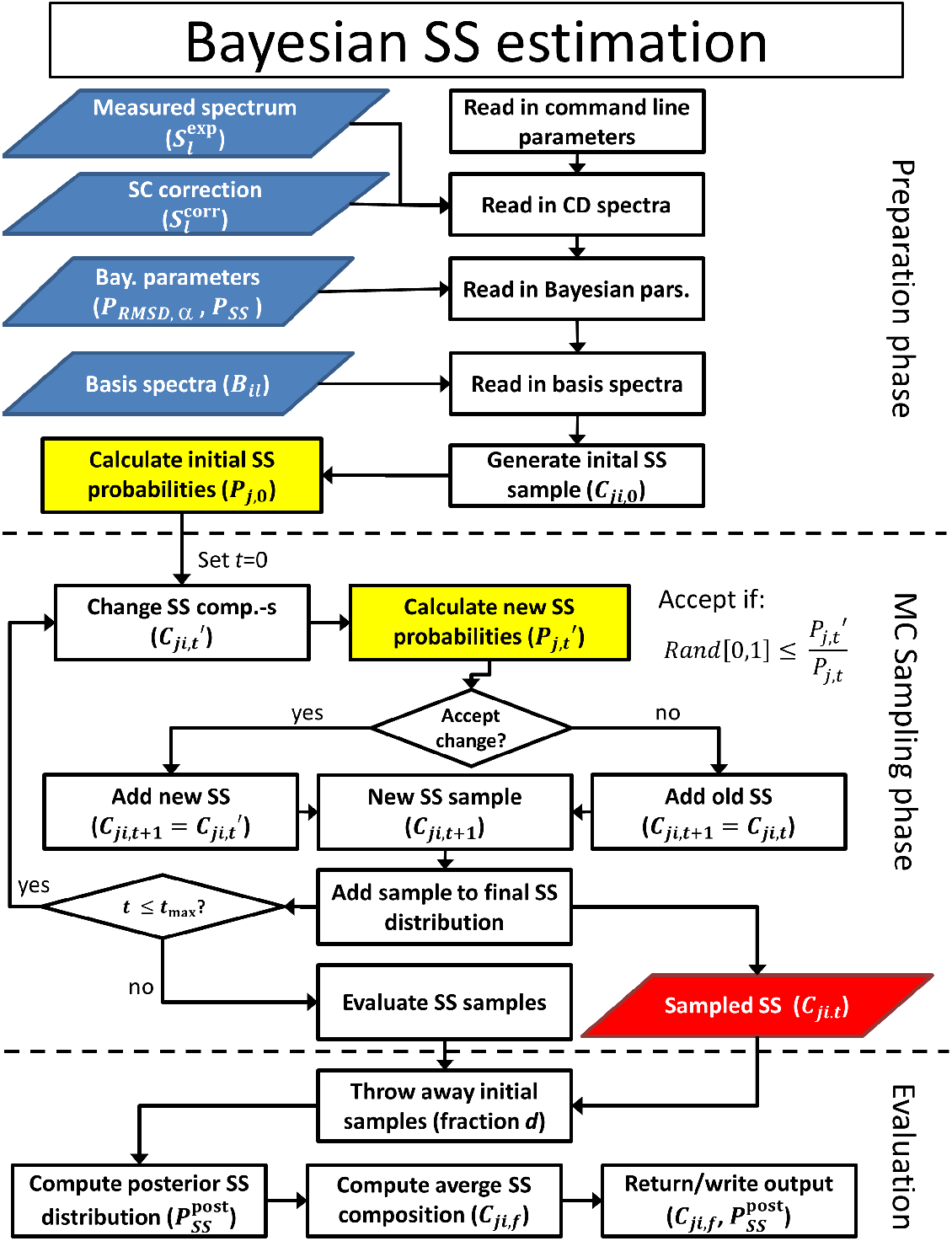
Schematic workflow of the Bayesian secondary structure estimation module in SESCA. The scheme depicts input data files as blue parallelograms, data on the sampled SS compositions are shown as a red parallelogram. Operations are depicted as white rectangles, and decisions are shown as white diamonds. Posterior probability calculation operations (see Fig. 1) are highlighted as yellow rectangles on the scheme.

### 4.1. Preparation and input parameters

In the preparation phase, input, output, and run parameters are read based on the user-provided command line arguments. If SESCA_bayes.py is used as a Python module, an array of arguments can be processed by the function *Read_Args* and passed to the *Main* function to run the algorithm. Arguments in SESCA are identified by preceding command flags (marked by the “@” character in the first position. There are four input files – shown as blue parallelograms in Fig. 3 – that SESCA_bayes accepts, each read in white-space separated data blocks stored as simple *ascii* text files.

The CD spectrum file (specified using the @spect flag) should contain two columns, wavelength in nanometers (nm) and CD signal intensity in 1000 mean residue ellipticity (kMRE) units. This file must be specified for SESCA_bayes, and if no command flags are provided, the first argument is automatically recognized as a CD spectrum file.

The side-chain correction file (specified by @corr) is an optional file to add baseline or sequence-dependent side-chain correction to the predicted CD spectrum, which are independent of the SS composition. If the basis spectrum set has basis spectra to calculate side-chain contributions, these signals can be computed before running SESCA_bayes, and added as a correction.

The Bayesian parameter file (@par) contains several data blocks, most importantly, the binned probability distribution function of CD deviations *P_RMSD,α_* (likelihood function), prior SS probability distributions for the SS composition *P* (*SS_j_*) and scaling factors *P* (*α_j_*), as well as the MC step parameters. If no parameter file is provided by the user, SESCA_bayes.py uses one of two default parameter files (Bayes_2D_SC.dat and Bayes_2D_noSC.dat) found in the “libs” sub-directory of SESCA, depending on whether a side chain correction file was provided or not. These files contain one of the two likelihood functions shown in Fig. 2, and uniform prior SS probability distributions. A more detailed description of the parameter blocks is provided in the examples sub-directory (examples 5).

The basis set file (@lib) contains several data blocks for CD spectrum calculations, including a block where the CD intensity of 3-6 basis spectra at each wavelength (175-269 nm) is provided. Several derived basis sets are available in libs sub-directory, and a detailed description of the data blocks is given in example 1.

In addition to the input files, SESCA bayes recognizes several additional command flags to modify program behavior. The number of initial SS compositions for MC sampling phase is specified by @size. The number of MC steps per initial SS composition is set by @iter. The @scale flag allows the user to control whether the measured CD spectrum is rescaled before determining the deviation from the predicted CD spectra or not. In the absence of these command flags, the values 100, 500, and 1 (yes) are used for the SS estimation.

Finally, three command flags control the output of SESCA_bayes.py; providing a “0” argument to any of these flags disables writing the associated output. The command flag @write specifies the file name for the primary output, and if no command flags are given, SESCA_bayes automatically recognizes the second argument as primary output file. This file contains a summary of the input parameters, binned posterior probability distributions for the SS compositions and scaling factors, as well as the most probable SS fractions and their uncertainties. The command flags @proj and @data allows the user to print secondary output files. The @proj flag specifies a file name for heatmap-style two-dimensional projection of the posterior SS distribution. The projection is made along two SS fractions selected using the @pdim flag Finally, the flag @data specifies a file name for printing all the sampled SS compositions the primary output is computed from, along with their estimated CD deviations, prior and posterior probabilities. By default, only the primary output file is printed into ‘SS_est.out’, and no secondary output is written.

### 4.2. Monte Carlo sampling

To determine the most probable SS composition of the protein based on its CD spectrum, sampling of the SS space is required. To this aim, SESCA_bayes uses a MC sampling scheme starting from N (set by @size) initial SS compositions, drawn from the prior SS distribution using rejection sampling. As the center part of Fig. 3 shows, at every step *t* of the MC sampling phase, a change on each of the SS compositions (*C_ji,t_*) is attempted. The change is realized by transferring a given SS fraction between two randomly chosen SS classes, yielding a new SS composition 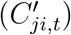. The amount of the transferred SS fraction from the donor class to the acceptor class is determined based on the distribution specified in the Bayesian parameter file. If no distribution is provided, the fraction is drawn from a Gaussian distribution with a mean of 0.05 and variance of 0.1. To remain in the space of possible SS compositions, the transferred SS fraction cannot exceed the current fraction assigned to the donor class, and classes that currently have a fraction of zero assigned to them cannot be selected as donors.

After the changes are attempted, the posterior probabilities 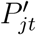 of the new SS compositions are calculated (see Section 2) and compared to the posterior probabilities (*P_jt_*) of the SS compositions before the change. The attempted change is accepted or rejected by applying the Metropolis criterion to the ratio of posterior probabilities, *i.e.* the change is accepted if the ratio 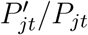 is larger than a randomly generated number between zero and one. If the change is accepted, 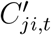 is added to the sampled SS distribution and used as the initial SS composition *C*_*ji,t*+1_ in the next MC step, otherwise *C_ji,t_* is added to the sampled SS distribution (again) and is used in the next MC step. This procedure is repeated until the specified number of MC attempts is reached, and yields *N × t_max_* sampled SS compositions. The sampled SS compositions resemble the prior SS distribution during the initial MC steps but converge towards an SS distribution weighted by the posterior SS probabilities.

### 4.3. Sample evaluation

The sampled SS distribution is analyzed in the evaluation phase, as shown in the bottom part of Fig 3. To avoid the over-representation of very low posterior probability SS compositions, a fraction of the initially sampled SS compositions may be discarded from final SS distribution. This fraction can be set by the user through the @discard flag, otherwise, the initial 5% of SS compositions is discarded. The remaining probability-weighted ensemble of possible SS compositions is used to compute the estimated SS composition 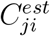 for the protein, the estimated scaling factor 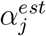, as well as to approximate the discrete posterior probability distribution for both quantities.

The estimated SS composition is determined by computing the mean and standard deviation (SD) of each SS fraction over the sampled SS compositions. Similarly, the most probable scaling factor is computed as the mean and SD of scaling factors estimated for the sampled SS compositions. The discrete probability distribution for both scaling factors and SS compositions are computed by binning all sampled SS compositions and scaling factors using the parameters extracted from the prior distributions provided in the Bayesian parameter file. The number of sampled SS compositions and scaling factors in each bin is normalized by the final sample size to obtain the discrete probability distributions. The computed estimates, their uncertainties and the discrete probability distributions are all written in the primary output file (defined by the @write flag) and returned as output by the SESCA_bayes module. If requested (@proj flag), the sampled SS compositions can be printed in a separate file. Finally, the two-dimensional projection of posterior SS distribution along two chosen SS fractions can also written into a separate output file (@proj flag), formatted as a human readable heat map, that can be easily processed into images using *e.g.* Python’s Matplotlib module [10] or external visualization programs.

## 5. Testing the Algorithm

### 5.1. Accuracy and precision

The accuracy and precision of the Bayesian SS estimation was tested using the 13 CD spectra listed in Table 1. Eight of these spectra (k= 1-8) are synthetic spectra that were generated from a given SS composition, but modified by adding artificial non-SS signals and scaling errors (see Section 3.2) to emulate CD deviations in real measured spectra. Four of the remaining five CD spectra (k= 9-12) are measured spectra from the SP175 set [8], for which the estimated SS compositions are compared to those extracted from the (protein data bank) structure of the reference protein. The last CD spectrum (k=13) was measured for the intrinsically disordered C-terminal domain of the Measles virus Nucleoprotein by Longhi *et al* [ref][11]. Because this domain is disordered, there was no experimental reference structure available for it, and therefore we used a molecular dynamics ensemble of Robustelli *et al* [9] as reference. This ensemble was generated using the Amber99SB-disp force field and was validated by small angle X-ray scattering (SAXS) and nucelar magnetic resonance (NMR) experiments. Table 1 lists the (estimated) CD deviations of all 13 CD spectra, quantified by the scaling factors *α_k_* and the root-mean-square intensity (*RMSI_k_*) of non-SS signals in each spectrum.

To test the accuracy of SESCA bayes, we estimated the SS composition of the above 13 CD spectra using the same DS-dT basis set with three SS classes (*α*-helix,*β*-strand, and Other) that was used to generate the synthetic spectra. The obtained Bayesian estimates for the test set are summarized in Table 2. This table includes the mean and SD (in parentheses) of SS fractions of the sampled posterior distributions, as well as the total SS deviation from the reference SS compositions, computed according to

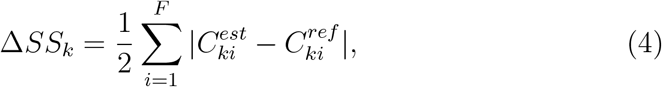

where 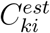 are the estimated SS fractions and 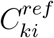 are the reference SS fractions listed in Table 1.

**Table 2:** Bayesian secondary structure estimates. The table lists the index and name of the model protein, the estimated fraction of its amino acids assigned to SS classes *α*-helix, *β*-strand, and Other, as well as the total SS deviation Δ*SS_k_* from the reference SS compositions shown in Table 1. The uncertainty (standard deviation) of each SS fraction and deviation is given in parentheses. Estimates that are more than 2 SD away from their reference value are highlighted in red.

| k | protein | α-helix | β-strand | Other | $\Delta SS_k$ |
| --- | --- | --- | --- | --- | --- |
| 1 | Synth-1 | 0.14 (0.05) | 0.44 (0.06) | 0.43 (0.04) | 0.06 (0.04) |
| 2 | Synth-2 | 0.49 (0.06) | 0.19 (0.06) | 0.32 (0.06) | 0.08 (0.05) |
| 3 | Synth-3 | 0.45 (0.06) | 0.12 (0.05) | 0.43 (0.05) | 0.03 (0.04) |
| 4 | Synth-4 | 0.22 (0.04) | 0.21 (0.05) | 0.57 (0.04) | 0.09 (0.04) |
| 5 | Synth-5 | 0.03 (0.04) | 0.26 (0.03) | 0.71 (0.05) | 0.07 (0.04) |
| 6 | Synth-6 | 0.36 (0.05) | 0.32 (0.04) | 0.31 (0.06) | 0.08 (0.04) |
| 7 | Synth-7 | 0.15 (0.04) | 0.01 (0.04) | 0.84 (0.05) | 0.05 (0.04) |
| 8 | Synth-8 | 0.03 (0.04) | 0.12 (0.04) | 0.85 (0.05) | 0.07 (0.04) |
| 9 | Lysm | 0.38 (0.05) | 0.04 (0.05) | 0.57 (0.05) | 0.05 (0.04) |
| 10 | Dqd-2 | 0.48 (0.06) | 0.06 (0.05) | 0.47 (0.05) | 0.12 (0.05) |
| 11 | Jacalin | 0.01 (0.04) | 0.31 (0.06) | 0.68 (0.06) | 0.03 (0.05) |
| 12 | Sub-C | 0.26 (0.05) | 0.13 (0.04) | 0.61 (0.04) | 0.04 (0.04) |
| 13 | MeV-Nt | 0.07 (0.05) | 0.13 (0.04) | 0.80 (0.05) | 0.10 (0.04) |

The obtained SS fractions show a fairly consistent 0.03 to 0.06 uncertainty. As expected, 35 of 39 SS fractions are within two SD of their reference value, with no significant difference in accuracy between synthetic and measured CD spectra, or globular and disordered protein models. In addition, the calculated total SS deviations (Δ*SS*) from the reference structures range between 0.03 and 0.12, and ten of thirteen values are also smaller than the estimated uncertainty of the estimation (two SD) that was calculated from the individual SD of SS fractions (*σ_ki_*) according to

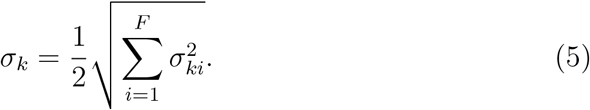

### 5.2. Comparison to deconvolution

Next, we compare the accuracy and precision of the Bayesian estimates to that of SS estimates obtained through spectrum deconvolution. To this aim, we estimated SS compositions with the deconvolution module of SESCA (SESCA_deconv) for the same 13 CD spectra (Table 1), using the same DS-dT basis spectrum set. The deconvolution was carried out using the most accurate protocol (method D2) tested previously [6]. This method constrains the basis spectrum coefficients to positive values, but normalizes them to unity only after the search for the best approximation. The SS compositions obtained using SESCA_deconv are listed in Table 3, along with the total SS deviations from reference SS compositions (found in Table 1). The total SS deviation of deconvolution estimates (Δ*SS_k_*) ranges from 0.0 to 0.29. The mean SS deviation for the whole set (0.07) is similar to that of the Bayesian estimates (0.07), but shows a significantly larger scatter (0.09 vs. 0.04). All three CD spectra with larger than average SS deviations (k= 3,4,10) have large non-SS contributions (2.0-2.9 kMRE), which is in line with our previous findings that non-SS contributions may be detrimental to the accuracy of deconvolution methods.

**Table 3:**
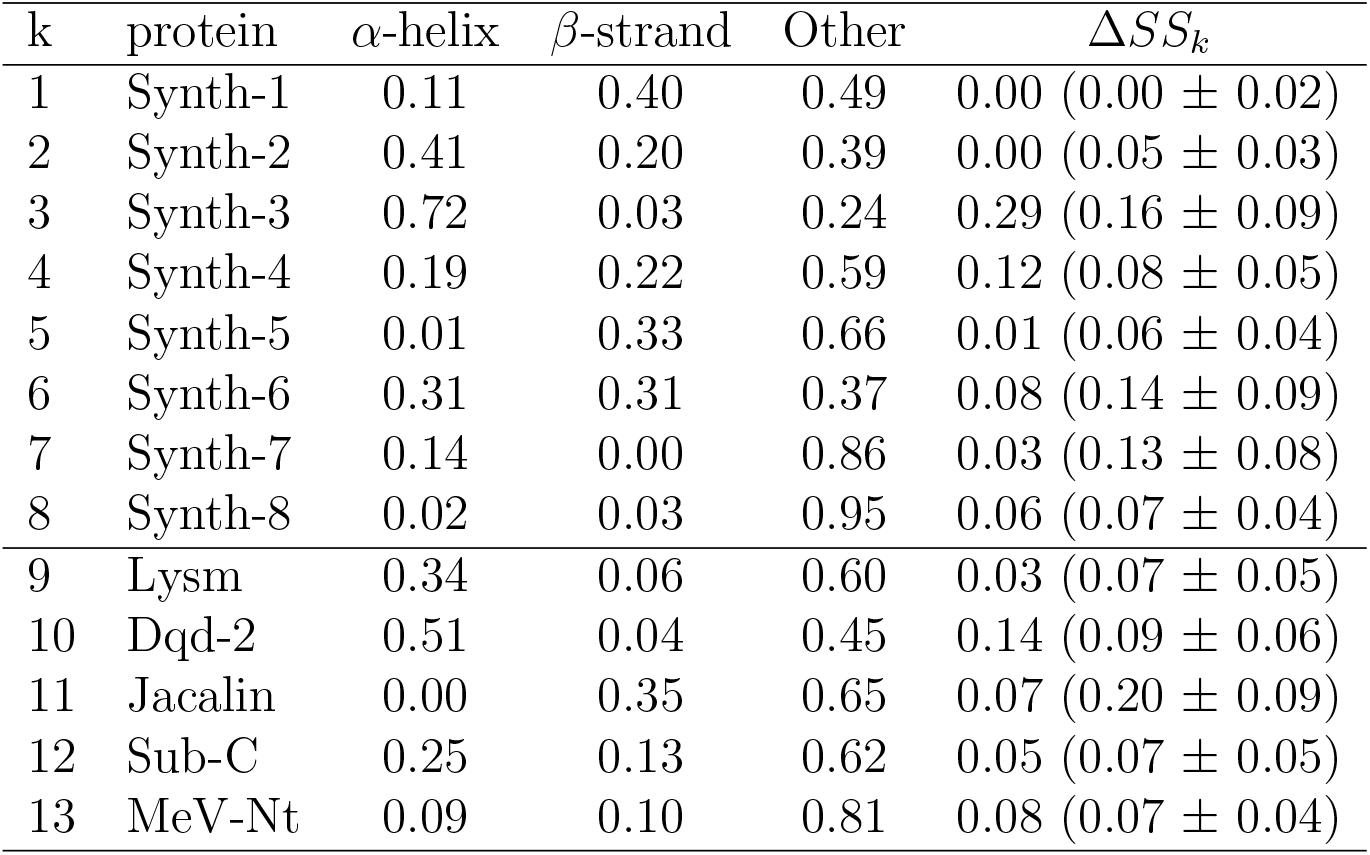
Secondary structure estimates based on spectrum deconvolution. The table lists the index and name of the model protein, the estimated fraction of its amino acids assigned to SS classes *α*-helix, *β*-strand, and Other, as well as the total SS deviation Δ*SS_k_* from the reference SS compositions shown in Table 1. The values in parentheses after Δ*SS_k_* show the mean and SD of the estimated total SS deviation computed from the rescaled RMSD between the measured (generated) CD spectrum and predicted spectrum of the SS estimate.

Although the SESCA deconv module does not provide information on the uncertainty of individual SS fractions, many SESCA basis sets (including DS-dT) include a calibration curve to estimate the expected total SS deviation if the true SS composition is unknown. This curve was computed from 4.9 × 10^5^ synthetic spectrum-structure combinations, which were binned according to their estimated non-SS contributions (*RMSD_j_*), to provide an expected mean and SD of SS deviations for a given (rescaled) RMSD. Comparing the true SS deviations of the deconvolution results with their estimated values shows that these estimates correctly describe the precision of the deconvolution method: 8 of 13 Δ*SS_k_* values are within 1 SD of the estimated total deviation, and all 13 fall within 2 SD. However, the average uncertainty of the deconvolution (0.09) is again considerably larger than that of the Bayesian SS estimates (0.04), and it increases with increasing non-SS contributions.

In summary, Bayesian SS estimation and spectrum deconvolution provides SS estimates that – in most cases – have a similar accuracy. However, Bayesian SS estimates are considerably more precise when significant non-SS contributions are present in the measured spectrum. Further, the Bayesian approach provides uncertainties for each individual SS fraction as well as for the optimal scaling factor of the measured CD spectrum, which is an additional advantage of the new method.

### 5.3. Example spectrum analysis

To further investigate the differences between the two methods, we analyzed the SS estimates for the CD spectrum with the largest difference between the deconvolution and Bayesian SS estimates. Figure 4A shows the obtained posterior SS probability distribution for synthetic spectrum 3, which contains larger than average non-SS contributions (2.02 kMRE). The heatmap shown in Fig. 4A illustrates that the most likely SS compositions are indeed clustered around the SS composition the synthetic spectrum was created from (shown as a red cross), with the highest posterior probability regions (shown in dark green) located in the immediate (Δ*SS_k_* < 0.05) vicinity of correct SS composition. However, the SS composition determined by deconvolution (purple cross) has a much higher *α*-helix content and it is not in a high-probability region in the Bayesian SS estimation.

**Figure 4:**
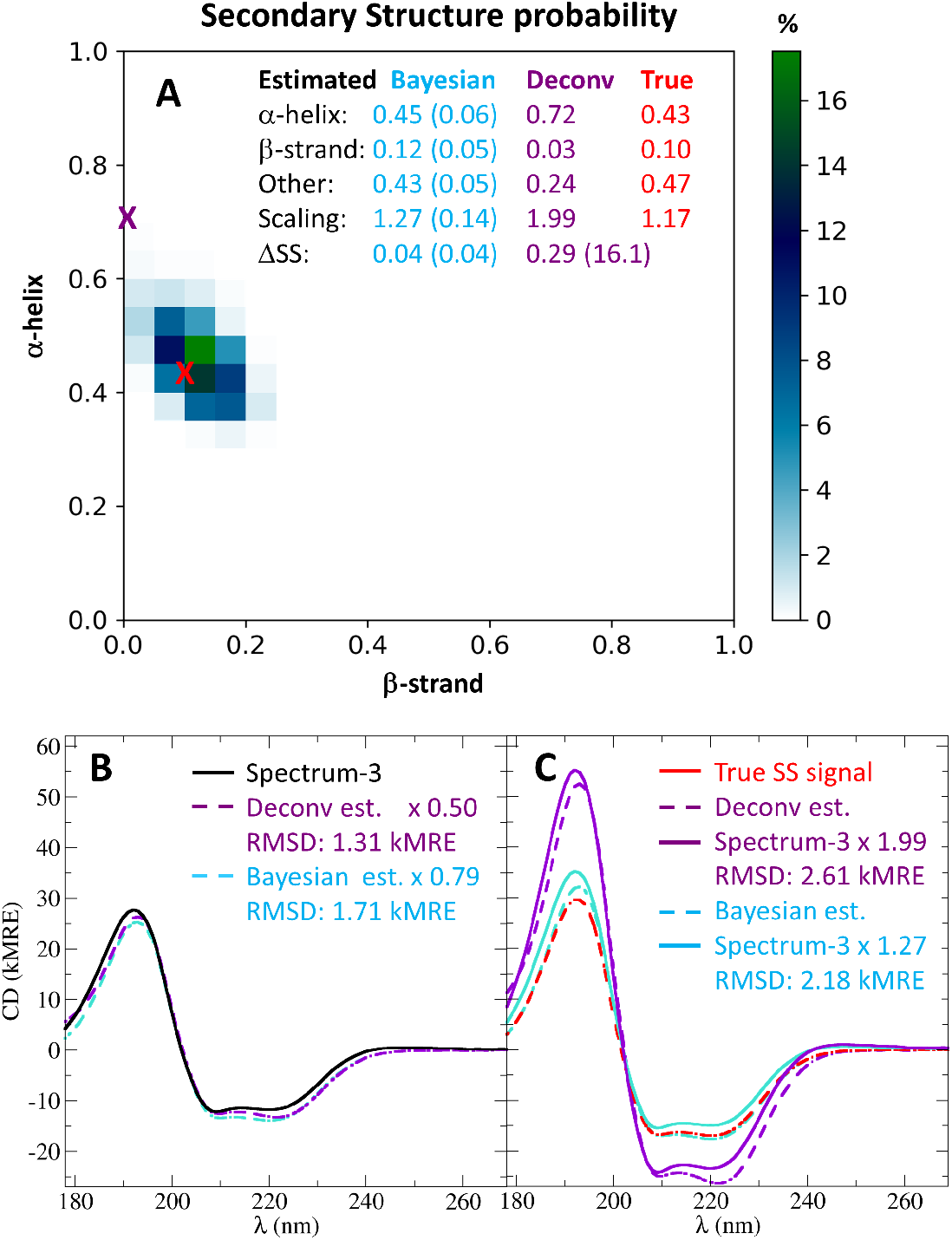
SS estimation for a synthetic CD spectrum. The figure compares the true SS composition (shown in red) with SS compositions obtained from Bayesian SS estimation (in blue) and spectrum deconvolution (in purple). A) shows the posterior probability distribution of sampled SS compositions in a heat map representation and indicates the true SS composition and the deconvolution SS estimate as crosses. The SS compositions, estimated scaling factors, and SS deviations are also listed in a tabulated format on the top. The difference on how the two estimates are evaluated by B) the deconvolution and C) Bayesian SS estimation is also shown. During deconvolution, the predicted CD signal of SS estimates is rescaled to match the measured CD spectrum, and the measure of quality is solely the RMSD. In the Bayesian approach, the measured spectrum is rescaled to match the predicted SS signals, and both the RMSD-s and the scaling factors are used to determine the most likely SS composition.

To examine why the two algorithms evaluate the proposed SS compositions differently, in Fig. 4B we computed the predicted CD signals of the two estimated SS compositions, rescaled them, and compared them to the synthetic spectrum, as is done during the deconvolution process. The figure shows that with the proper scaling factor both SS compositions approximate the synthetic spectrum well, but the deconvolution estimate (purple dashed line, *RMSD_j_* : 1.31 kMRE) fits slightly better than the Bayesian estimate (blue dashed lines, *RMSD_j_* : 1.71 kMRE).

In contrast, the Bayesian SS estimation rescales the synthetic CD spectrum to match the predicted spectra, and evaluates the likelihood of the SS compositions based on the joint probability of their non-SS contributions *RMSD_j_* and their scaling factors *α_j_*, as shown in Fig. 4C. Although the two estimates have a comparable RMSD in this method as well, the deconvolution estimate requires a scaling factor (*α_j_*: 1.99) to achieve a good agreement that is shown to be very unlikely according to the joint-probability map in Fig. 2A. Comparing the two estimated SS signals (dashed lines) to the SS signal of the true SS composition (in red) illustrates how considering scaling factors improves the precision of the SESCA_bayes. In this case, eliminating SS compositions with unlikely scaling factors from the sampled distribution allowed a fairly accurate (RMSD: 0.99 kMRE) approximation of the true SS signal.

## Appendix A. Declarations

## Acknowledgements

The authors would like to thank K. Blom and P. Kellers for the aid in editing the manuscript, as well as S. Longhi and S. Piana for providing the measured CD spectrum and molecular dnyamics ensemble of MeV-Nt, respectively.

## Authors’ contributions

G.N. designed and performed the computational analysis, and implemented code improvements. H.G. is the corresponding author, and assisted with the conceptualization. Both authors contributed to writing the manuscript.

## License

SESCA is free available under GNU general public license 3 (GPLv3).

## Code availability

the new SESCA implementation based on this study is available at: https://www.mpibpc.mpg.de/sesca

## References

[1] G. D. Fasman (Ed.), Circular Dichroism and the Conformational Analysis of Biomolecules, Springer US, Boston, MA, 1996 (1996). URL http://link.springer.com/10.1007/978-1-4757-2508-7

[2] N. J. Greenfield, Methods to estimate the conformation of proteins and polypeptides from circular dichroism data, Analytical biochemistry 235 (1) (1996) 1–10 (1996).

[3] S. M. Kelly, T. J. Jess, N. C. Price, How to study proteins by circular dichroism, Biochimica et Biophysica Acta (BBA) - Proteins and Proteomics 1751 (2) (2005) 119–139 (Aug. 2005). doi:10.1016/j.bbapap.2005.06.005. URL http://linkinghub.elsevier.com/retrieve/pii/S1570963905001792

[4] S. Brahms, J. Brahms, Determination of Protein Secondary Structure in Solution by Vacuum Ultraviolet Circular Dichroism, Journal of Molecular Biology 138 (1980) 147–178 (1980).

[5] G. Nagy, M. Igaev, N. C. Jones, S. V. Hoffmann, H. Grubmüller, SESCA : Predicting Circular Dichroism Spectra from Protein Molecular Structures, Journal of Chemical Theory and Computation (Aug. 2019). doi:10.1021/acs.jctc.9b00203. URL http://pubs.acs.org/doi/10.1021/acs.jctc.9b00203

[6] G. Nagy, H. Grubmüller, How accurate is circular dichroism-based model validation?, European Biophysics Journal 49 (6) (2020) 497–510 (Sep. 2020). doi:10.1007/s00249-020-01457-6. URL http://link.springer.com/10.1007/s00249-020-01457-6

[7] A. Gelman, J. B. Carlin, Bayesian Data Analysis, 3rd Edition, Texts in Statistical Science, Chapman and Hall/CRC, 2014 (2014).

[8] J. G. Lees, A. J. Miles, F. Wien, B. A. Wallace, A reference database for circular dichroism spectroscopy covering fold and secondary structure space, Bioinformatics 22 (16) (2006) 1955–1962 (Aug. 2006). doi:10.1093/bioinformatics/btl327. URL http://bioinformatics.oxfordjournals.org/cgi/doi/10.1093/bioinformatics/btl327

[9] P. Robustelli, S. Piana, D. E. Shaw, Developing a molecular dynamics force field for both folded and disordered protein states, Proceedings of the National Academy of Sciences 115 (21) (2018) E4758–E4766 (2018).

[10] J. D. Hunter, Matplotlib: A 2d graphics environment, Computing in Science & Engineering 9 (3) (2007) 90–95 (2007). doi:10.1109/MCSE.2007.55.

[11] S. Longhi, V. Receveur-Bréchot, D. Karlin, K. Johansson, H. Darbon, D. Bhella, R. Yeo, S. Finet, B. Canard, The C-terminal Domain of the Measles Virus Nucleoprotein Is Intrinsically Disordered and Folds upon Binding to the C-terminal Moiety of the Phosphoprotein, Journal of Biological Chemistry 278 (20) (2003) 18638–18648 (May 2003). doi:10.1074/jbc.M300518200. URL https://linkinghub.elsevier.com/retrieve/pii/S0021925819549730

